# Predicting and Elucidating Peptide Retention Mechanisms with Graph Attention Networks

**DOI:** 10.64898/2026.05.18.725893

**Authors:** Alexander Kensert, Katerina Hruzova, Robbe Devreese, Alireza Nameni, Arthur Declercq, Ralf Gabriels, Lennart Martens, Robbin Bouwmeester, Jiri Urban

## Abstract

Liquid chromatography (LC) is a key technology in bottom-up proteomics, separating proteolytic peptides to decrease sample complexity, enhance coverage, and increase the robustness of protein identification and quantification. Although high-resolution mass spectrometry has advanced significantly, comparable progress in LC has lagged, primarily due to a limited understanding of peptide-column interactions. To bridge this knowledge gap, we introduce a novel deep learning model (PeptideGNN) based on a Graph Neural Network (GNN) architecture to model and elucidate peptide behaviors across various separation conditions. Trained to accurately predict peptide retention times on ten diverse proteomic datasets, the model subsequently employed a saliency mapping technique to interpret the underlying retention mechanisms. Our model consistently outperformed existing retention-time predictors across multiple datasets, while the saliency mapping, importantly, revealed insights into peptide-stationary phase interactions, highlighting the effects of neighboring amino acids, post-translational modifications (PTMs), chromatographic columns, and mobile phase additives on peptide retention.

## Introduction

Conventionally, peptides are encoded as character sequences, where each character represents a specific amino acid residue linked by peptide bonds^1^. Deep learning architectures, such as Recurrent Neural Networks (RNNs) or Convolutional Neural Networks (CNNs), utilize these sequences to project peptides into a latent embedding space optimized for specific predictive tasks. However, a significant limitation of these sequence-based models is their reliance on coarse-grained encodings which only capture the sequence order of amino acids. Consequently, they fail to generalize to amino acid modifications or capture the atomic and substructural patterns essential for understanding complex phenomena like chromatographic retention. To address this deficiency, we introduce a novel hybrid architecture that integrates Graph Neural Networks (GNNs) with RNNs. This approach allows us to simultaneously capture fine-grained atomic interactions and long-range sequential dependencies (**Fig. 1**). By capturing this molecular detail, our hybrid approach aims to overcome the limitations of conventional sequence-based models, offering a path toward improved prediction accuracy and clearer insight into peptide retention mechanisms.

**Figure 1.**
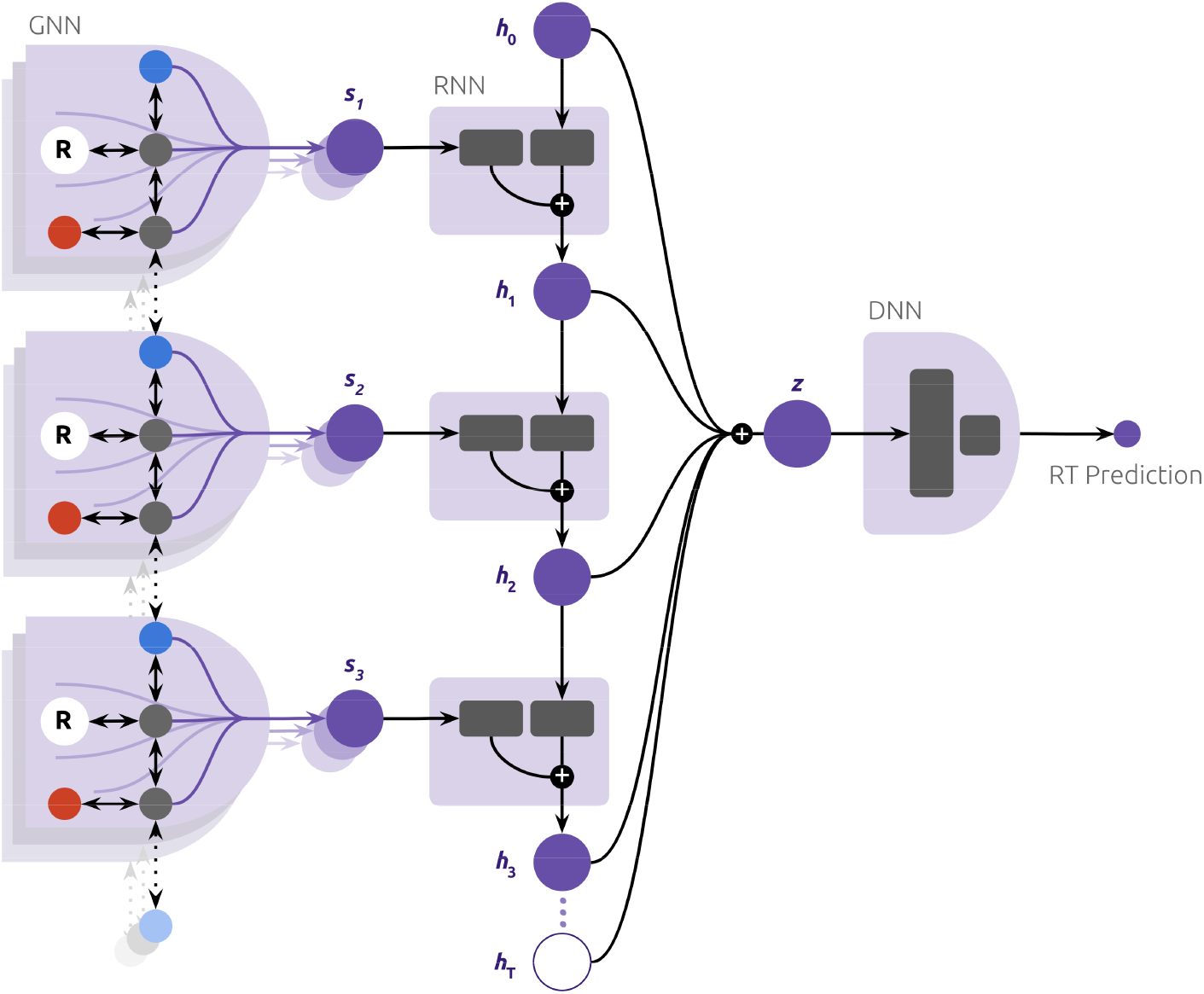
A high-level and simplified illustration of the model architecture of this study. The figure illustrates the combination of a GNN, RNN and DNN architecture to model peptides in the context of chromatography. Important architectural components not shown in the figure: attention heads and bidirectionality of the RNN. The symbol R denotes the sidechain subgraph, which is compressed into a single node for improved visualization; the symbols s_t_ denote the learned amino acid embeddings; the symbols h_t_ denote the learned (hidden) states along the peptide sequence; and the symbol z denotes the peptide embedding; and the RT prediction denotes the retention time prediction.

While peptide modeling for chromatography and mass spectrometry dates back to the 1980s, primarily utilizing amino acid composition^2,3^, highly accurate deep learning models have a much more recent origin, beginning in the late 2010s. The first notable example came in 2018 from Ma et al.^4^, who implemented a capsule CNN to predict retention times with substantially greater accuracy compared to earlier classical machine learning models. Following this, in 2019, the application of RNNs gained traction, with Guan et al.^5^ using the architecture to predict various LC-MS/MS peptide behaviors, including retention time, and Gessulat et al.^6^ employing an RNN for the prediction of retention times and peptide fragment intensities. The trend toward hybrid models began in 2020 when Yang et al.^7^ combined a CNN and an RNN to predict both RT and fragmentation intensities. A significant step toward handling modifications was taken in 2021 by Bouwmeester et al.^8^ with DeepLC, which used CNNs and explicitly encoded atoms to predict RTs for modified peptides, even those unseen during training. More recently, between 2021 and 2023, several studies utilized the Transformer architecture to model peptides for various predictions, including RT, fragmentation, and other property predictions^9–12^. However, a study by Franklin et al.^13^ observed a decline in performance when using transformers for peptide modeling (specifically, RT and collisional cross section predictions) compared to CNNs and RNNs.

Despite substantial progress in peptide modeling for chromatography and mass spectrometry, current state-of-the-art approaches largely overlook low-level structural information and model interpretability. To address this critical gap, we here introduce a novel framework that explicitly models peptides as molecular graphs, defining atoms as nodes and chemical bonds as edges. By leveraging this fine-grained graph representation, our hybrid GNN-RNN architecture not only achieves highly accurate and precise retention time predictions but also enables researchers to elucidate the complex and often obscure mechanisms governing chromatographic retention. The subsequent sections detail the architecture, model training, performance evaluation against existing predictors, and the mechanistic insights derived from our interpretability analysis.

## Methods

### Peptide graph

As a large molecule, a peptide can naturally be represented as a graph *G = (V, E, X, F)*, where *V =*{*v*_*1*_, *v*_*2*_,*…, v*_*3*_} represents a set of nodes corresponding to atoms, *E =* {*e*_1_, *e*_2_,…, *e*_*m*_} represents a set of edges corresponding to bonds, *X =* {*x*_*1*_, *x*_*2*_, *…, x*_*n*_} denotes a set of node feature vectors corresponding to the nodes in *V*, and *F =* {*f*_1_, *f*_*2*_, *…, f*_m_*}* denotes a set of edge feature vectors corresponding to the edges in *E*; *n* and *m* denotes the number of nodes and edges of the peptide graph, respectively. If the edge represents the edge between nodes _*i*_ and _*j*_, the edge is denoted *e*_*ij*_ and its feature vector is denoted *f*_*ij*_. Although the bonds in a peptide are undirected, the edges of the peptide graph were, in this study, bidirectional; namely, for every edge *(i, j) ∈ E* the reverse edge *(j, i) ∈ E* also existed, and their feature vectors, *f*_*ij*_ and *f*_*ji*_, while encoded in separate vectors, equaled each other. This representation is similar to the representations previously used for small molecules^14^.

In this study, and in the context of molecular graphs, the feature vectors *X* and *F* encoded atom and bond features, such as atom and bond type, respectively. **Table 1** summarizes the features used in this study, which were all computed using the open-source cheminformatics software RDKit^15^. For more information regarding the features, we refer to RDKit’s documentation^16^.

**Table 1.** Atom and bond features used in this study.

| Feature type | Feature set | Feature encoding |
| --- | --- | --- |
| **Atom** |  |  |
| Atom type | C, N, O, P, and S | One-hot |
| Hybridization | s, sp, sp2, sp3, sp3d, sp3d2, unspecified | One-hot |
| Formal charge | -3, -2, -1, 0, 1, 2, 3, and 4 | One-hot |
| Chirality | R, S, None | One-hot |
| Degree | 0, 1, 2, 3, 4, 5, 6, 7, and 8 | One-hot |
| Number of valences | 0, 1, 2, 3, 4, 5, 6, 7, and 8 | One-hot |
| Number of hydrogens | 0, 1, 2, 3, and 4 | One-hot |
| Part of ring of size n | 0, 3, 4, 5, 6, 7, and 8 | One-hot |
| Number of radical electrons | 0, 1, 2, 3 | One-hot |
| Aromatic |  | Binary |
| Chiral center |  | Binary |
| Heteroatom |  | Binary |
| Hydrogen donor |  | Binary |
| Hydrogen acceptor |  | Binary |
| Part of ring |  | Binary |
| Gasteiger charge |  | Continuous |
| Log P contribution |  | Continuous |
| Total polar surface area contribution |  | Continuous |
| **Bond** |  |  |
| Bond Type | Single, Double, Triple, Aromatic | One-hot |
| Conjugated |  | Binary |
| Stereo | E, Z, None | One-hot |
| Rotatable |  | Binary |

### Graph Neural Network

The GNN architecture employs *L = 3* layers, each integrated with *K = 8* independent attention heads to enhance representational learning^17,18^. In each layer, node features undergo a learned transformation (Eq. 1) before being aggregated with those of their nearest neighbors (Eq. 2). To capture the relative significance of local substructures, the aggregation step utilizes an attention mechanism that dynamically weighs the influence of neighboring nodes during the update process (Eq. 3 and 4).

Formally, the first step, the *transformation*, is defined as follows:

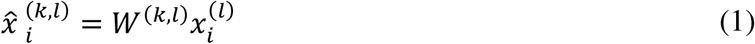

where 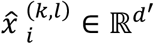 denotes transformed node feature vector of *v*_*i*_ for layer *l* and attention head *k*, with 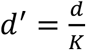 *and d = 128* ; and 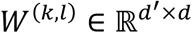 denotes the learnable weight matrix for layer *l* and attention head *k*. Notably, prior to the first GNN layer, atom and bond features *d = 128* re projected, via learnable linear transformations, to match the dimension of *d = 128*, ensuring consistent dimensionality across GNN layers.

The second step, the *aggregation*, is defined as follows:

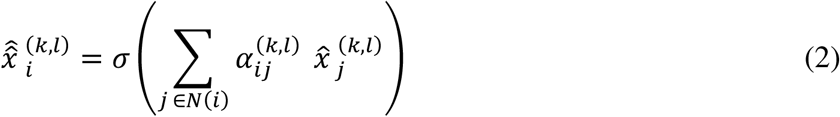

where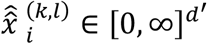 denotes the aggregated node feature vector, encoding information up to radius *l* of node *v*_*i*_; *a* denotes a non-linear activation function, ReLU; and 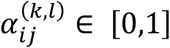 denotes the normalized attention coefficient, computed via a Softmax function as follows:

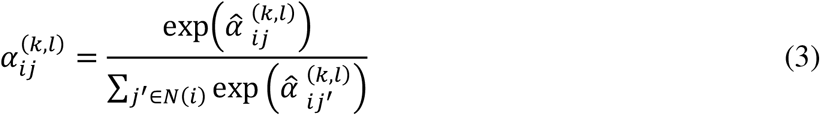

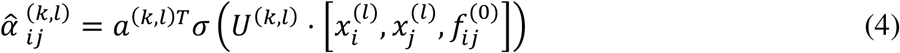

where *U*^*(k,l)*^ ∈ ℝ^*d×3d*^ and *a*^*(k,l)*^ ∈ ℝ^*d*^ denote a learnable weight matrix and a learnable weight vector, respectively. This learnable weight matrix allows the GNN to selectively focus, or attend, on specific nodes or substructures within the graph. *a* denotes another non-linear activation function, LeakyReLU.

After the transformation and aggregation, the GNN layer combines the attention heads by concatenating their outputs as follows:

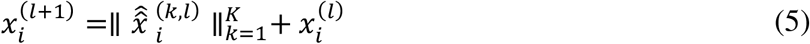

where 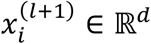 denotes the updated note feature vector. 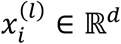 denotes the (previous) node feature vector, added to preserve information and prevent vanishing gradients^19^.

After *L* layers of transformations and aggregations, the resulting node features 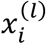 of each layer *l*, including the initial node features 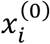, were concatenated to produce the final node embedding as follows:

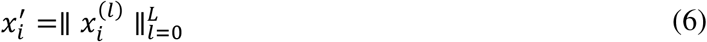

where 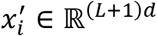 denotes the embedding for node *v*_*i*_. Notably, 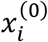 and 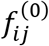 represent the linearly transformed atom and bond feature vector for node *v*_*i*_ and edge *e*_*ij*_, respectively.

### Recurrent Neural Network

The recurrent neural network (RNN) of this study was a long-short term memory (LSTM)^20^ model, which allowed the peptide model to capture high-level interactions within the peptide. To reduce node embeddings to subgraph (amino acid) embeddings, a local average pooling operation was performed on the node embeddings as follows:

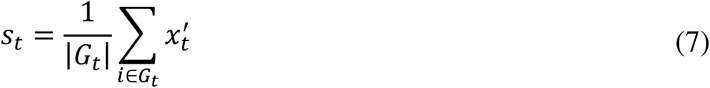

where *s*_*t*_ ∈ ℝ^(*L+)d*^ denotes the t:th amino acid embedding of the peptide sequence. The resulting sequence of amino acid embeddings, *S =* {*s*_*1*_, *…, s*_*T*_}, where *T* denotes the number of amino acids in the peptide, was sequentially passed to the LSTM as input.

The LSTM maintained a cell state *c*_*t*_ ∈ ℝ^*d*^ and a hidden state *h*_*t*_ ∈ ℝ^*d*^ over the input sequence {*s*_*1*_, *…s*_*T*_}, which served as its long-term memory and short-term memory, respectively; allowing the LSTM to both model longer-distance interactions, such as electrostatic interactions from folding and disulfide bonds, as well as shorter-distance interactions, such as hydrogen bonds formed between backbone atoms or side-chain atoms in close proximity, respectively.

Specifically, the LSTM utilized three different gates that decided what information from the previous cell state *c*_*t-1*_ should be forgotten, what new information from the input *s*_*t*_ and previous hidden state *h*_*t-1*_ should be stored in the new cell state *c*_*t*_, and what part of the new cell state *c*_*t*_ should be used to compute the new hidden state *h*_*t*_ :

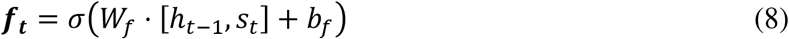

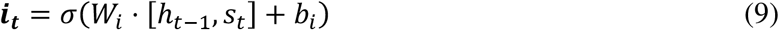

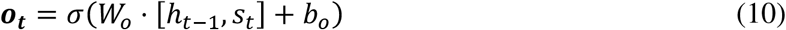

where ***f***_*t*_ ∈ [0,1]^*d*^, ***i***_*t*_ ∈ [0,1]^*d*^, and *o*_*t*_ *∈ 0,1]*^*d*^, denote the forget gate, input gate and output gate, respectively. *a* is another non-linear activation function, specifically a sigmoidal function; and *W*_*f*_, *W*_*i*_, *W*_*o*_, *b*_*f*_, *b*_*i*_ and *b*_*o*_ are learnable weight matrices and bias vectors.

Before computing the cell state *c*_*t*_, a candidate cell state 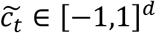 was computed as follows:

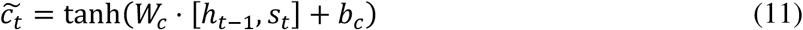

where *W*_*c*_ and *b*_*c*_ is the learnable weight matrix and bias vector, respectively. The cell state was then updated by combining the previous cell state *c*_*t-l*_ and the candidate cell state 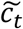, scaled by the forget gate ***f***_*t*_ and input gate ***i***_*t*_, respectively:

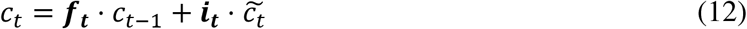

The hidden state was then computed using the output gate ***o***_*t*_, as follows:

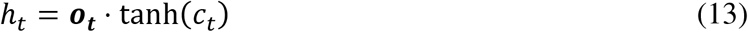

After *T* steps, the hidden states {*h*_1_, *…, h*_*T*_}, were averaged to obtain the peptide embedding *z* ∈ ℝ^*d*^, which was subsequently passed to a prediction network to produce an RT prediction for the peptide. As the peptide sequence has no real direction (unlike a piece of text, which has a direction), the LSTM propagated information from the C-terminal to N-terminal, as well as from the N-terminal to the C-terminal, resulting in *z* ∈ ℝ^*d*^. This implementation is often referred to as a bidirectional LSTM.

### Dense Neural Network

The prediction network of this study was a Dense Neural Network (DNN), often referred to as a Multi-Layer Perceptron (MLP), configured to predict RT. The model consisted of two layers in total, one hidden layer with 1024 units and a ReLU activation function, and one output layer with a single unit for the RT prediction. Formally, the RT prediction *ŷ* is mapped from the peptide embedding *z* through successive fully connected transformations, as follows:

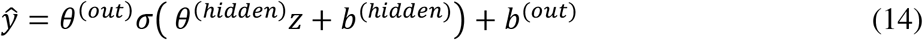

where *θ*^*(hidden)*^, *θ*^*(out)*^, *b*^*(hidde)*^, *b*^*(out)*^are the learnable weight matrices and bias vectors, and *a* the ReLU activation.

### Saliency mapping for interpretability

For interpretability and elucidation of retention mechanisms, saliency mapping based on Grad-CAM^21,22^ was implemented. In brief, the saliency mapping first performed the forward pass (Eq. X, Y, Z), then computed the partial derivatives of the output with respect to the inputs (specifically, the node features, at every layer). Specifically, scalar saliency values ψ_*i*_ for each node *v*_*i*_ were obtained as follows:

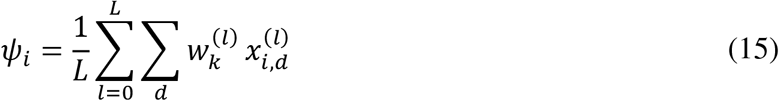

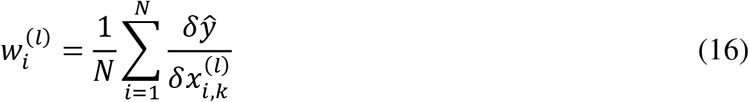

where 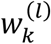 denotes the importance weight for feature *k* after layer *l*. The value in 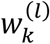 defines the importance of a certain feature *k*. A larger absolute value of 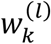 indicates a higher importance for that feature. The sign of the weight shows its contribution: a positive value means the feature positively influences the RT, while a negative value indicates a negative influence. For example, if 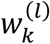 has a large positive value, and a certain node encodes this feature, then this node, considering this specific feature, contributes positively to RT. *w*_*i*_ therefore gives us the total contribution of the node to the RT, and the contribution may be positive or negative. We refer to positive contributions of *w*i in the saliency map as molecular retention, while negative contributions promote repulsion. A similar approach has been used previously when solubility–insolubility or toxicity–non–toxicity from molecular feature maps were predicted and interpreted^23^.

Furthermore, depending on the desired resolution of the retention mechanism interpretation, either atom or amino acid saliencies were obtained, either by directly using the saliency values of ψ_*i*_ or by performing a local sum pooling as follows:

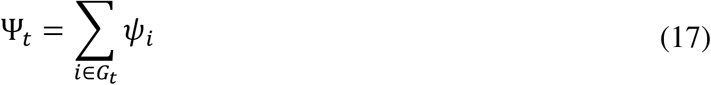

where Ψ_*t*_ denotes the scalar saliency value of subgraph (amino acid) *t* in the peptide sequence.

The magnitude of the saliency values depends on both the magnitude of the retention times and the chromatographic setup being studied. To preserve all relevant information, saliency values were used unnormalized. However, to visualize the magnitude of peptide interactions across different chromatographic setups (**Fig. 3b, 3c**, and **Fig. 6c**), normalization was performed at both the dataset and peptide levels. For this normalization, each selected dataset was normalized by dividing saliency values by the maximum absolute value within that dataset. Subsequently, the peptide common to all datasets was selected and further normalized by dividing the saliency values by the maximum absolute value observed for that peptide across datasets.

### Datasets

Ten datasets with various chromatographic conditions were investigated, including five reversed-phase (RP) datasets, four hydrophilic interaction (HILIC) datasets, and one cation-exchange (SCX) dataset. DIA HF^24^, SWATH Library^25^, and SCX^26^ were selected as they are common benchmarks for RT prediction models. Three smaller reversed-phase datasets FA^27^, AcA^27^, and EG^28^, were included to investigate the dataset size effect and the influence of mobile phase additives, including formic acid (FA), acetic acid (AcA), and the addition of ethylene glycol (EG) to the acidified mobile phase. Samples from the same laboratory, determined under identical experimental conditions, were obtained to study the retention for different HILIC columns (LUNA HILIC, LUNA SILICA, ATLANTIS SILICA, XBridge)^29^. Table SI-1 in the Supporting Information provides detailed information about the experimental parameters used for each dataset. DIA HF and SWATH Library datasets used indexed retention time (iRT)^30^.

Each dataset was randomly split into a training set (70%), a test set (15%), and a validation set (15%). For the performance comparison with the DeepLC model^8^, datasets were split accordingly (85% for the training set, 10% for the test set, and 5% for the validation set). Performance results were computed on a test set, and the evaluation metrics included Pearson correlation (R), mean absolute error (MAE), and Δt_95%_, which represents the error in retention time prediction within a 95% confidence interval.

Datasets FA, AcA, and EG were downloaded from the jPOST repository^31^ and the PRIDE repository^32^ and raw MS files were processed by MaxQuant^33^ with FDR < 0.01. Oxidation of methionine and N-terminal acetylation were set as variable modifications, and carbamidomethylation of cysteine was set as a fixed modification. The identifications were filtered on posterior error probability ≤ 0.001. The MaxQuant evidence files were processed using the ProteomicsML protocol for retention time alignment^34^. Peptidoform retention times were aligned across runs using splines in a Generalized Additive Model implemented via pyGAM LinearGAM. An initial reference run was selected based on the minimal number of missing peptidoforms. Each run was then iteratively aligned to the reference run, which was updated as the median of the aligned runs, enabling refinement of fitting. Datasets FA and AcA were split based on the mobile phase additive and the laboratory of origin before the retention time alignment.

## Results

### Predictive performance

The prediction accuracy of PeptideGNN is compared to DeepLC on a set of diverse datasets (**Fig. 2**). Across all datasets, PeptideGNN achieved, on average, a 10% lower MAE value, providing consistent improvement. For completeness, the scatter plots of predicted versus observed retention times for both models are provided in the Supporting Information (**Fig. SI-1**).

**Figure 2.**
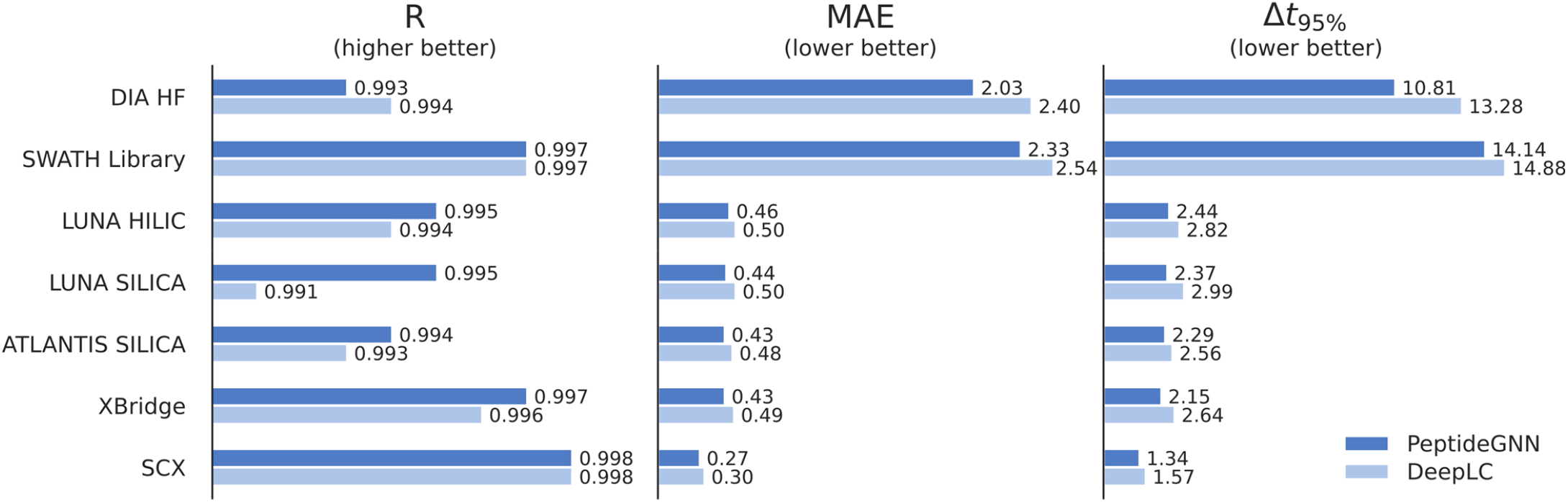
Comparison of PeptideGNN model performance with DeepLC on seven datasets. The performance was compared on test sets for various datasets using correlation coefficient (R), mean absolute error (MAE, min), and the error in retention time prediction within a 95% confidence interval (Δt_95%_, min). Retention times for the DIA HF and SWATH Library datasets were expressed as indexed retention times.

**Figure 3.**
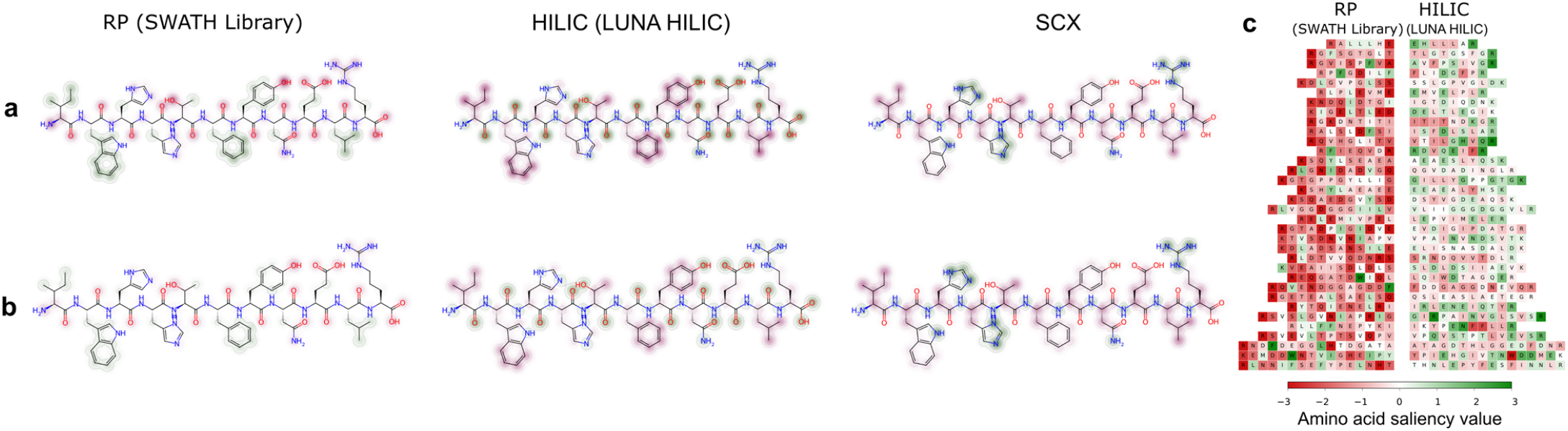
Peptide saliency patterns under three retention mechanisms, shown as (a) unnormalized, and (b) normalized saliency maps; (c) shows a comparison chart of amino acid contributions for identical peptides (displayed as mirrored sequences) found in two retention mechanisms. The green color indicates a positive contribution to retention, whereas the red color indicates a negative contribution to retention. The peptide depicted in (a) and (b) is IWHHTFYNELR. In (b) and (c), both dataset normalization and peptide normalization were performed on atomic saliency values.

### Retention visualized by saliency mapping

To evaluate the interpretability of PeptideGNN, we generated saliency maps that highlight regions in the molecule that contribute most to retention time prediction. When the analyte has no affinity to the stationary phase, it passes through the column with the mobile phase velocity, and the resulting retention time is equal to the void time of the column. As the affinity of the analyte to the stationary phase increases, the retention time increases as well.

**Fig. 3** presents unnormalized (**Fig. 3a**) and normalized (**Fig. 3b**) saliency maps for the peptide IWHHTFYNELR across RP, HILIC, and SCX datasets and allows the comparison of individual retention mechanisms. In the unnormalized maps (**Fig. 3a**), each molecule is scaled independently: the atom with the largest absolute effect on RT prediction is assigned to the maximum number of contour rings, and the remaining atoms are scaled accordingly. In contrast, the normalized maps (**Fig. 3b**) apply the scaling across datasets for the selected peptide. The saliency values were first normalized at the dataset level by dividing by the maximum absolute saliency value, and subsequently at the peptide level by assigning the maximum number of contour rings to the atom found across selected molecules. While the unnormalized maps provide the most straightforward visualization for interpreting a single dataset, the normalized maps are suitable for comparing interaction strengths across different models or chromatographic conditions.

The raw saliency maps (**Fig. 3a**) show that, in RP, the highest positive saliency (green color in the map) is found for nonpolar residues (phenylalanine, leucine, isoleucine), while negative saliency (red color in the map) is found for polar residues (glutamine, asparagine). These saliency maps reflect the fundamental principles of RP chromatography, in which retention is predominantly governed by hydrophobic partitioning: aliphatic and aromatic residues promote interaction with the stationary phase and thus retention, whereas polar residues interact more strongly with the mobile phase and therefore reduce retention^35^.

In the HILIC dataset, the same peptide exhibits the opposite pattern: charged residues (histidine and glutamate) show strong positive saliency, whereas aliphatic and aromatic residues (phenylalanine, leucine, and isoleucine) have a negative contribution. These patterns reflect the expected Coulombic interactions between charged residues and the aqueous layer, which is immobilized on the polar surface of the stationary phase, while hydrophobic residues remain preferentially in the organic-rich mobile phase^35^.

SCX is based on the strong electrostatic interaction between the negatively charged stationary phase and the positively charged analyte^26^. The saliency map for the SCX dataset confirms this principle: only positively charged residues (histidine and arginine) show positive contributions to retention. Whereas negatively charged and nonpolar residues either reduce retention or have a negligible contribution.

For this peptide sequence, the normalized saliency maps (**Fig. 3b**) indicate the strongest interaction under the SCX mechanism and the weakest interaction under the RP mechanism. This observation correlates with the peptide retention time in selected datasets, as the presented peptide exhibits high retention in SCX and low retention in RP.

To further investigate the retention mechanism patterns, saliency maps were generated for peptides shared across all ten datasets (**Fig. SI-2** to **SI-8**). Across datasets, the maps consistently reproduce the expected retention mechanisms. This demonstrates that PeptideGNN is able to model the underlying retention logic directly from the peptide sequence and associated retention times.

### Amino acid contributions on the peptide level

Saliency patterns at the molecular level generally reflect retention trends observed in small molecule separations^36^, but the unique physicochemical properties of peptides require a different approach for their separation in liquid chromatography. Tryptic peptides are polyprotic ampholytes with varying sizes, chelating properties, and are inherently chemically reactive. This, for example, leads to the protonation of the N-terminal amino group and basic residues in the typically acidic mobile phase of LC-MS experiments. Furthermore, depending on the mobile phase pH, the carboxyl group on the C-terminal is typically uncharged or deprotonated. The side-chain contributions of each amino acid to retention depend significantly on its pKa, which is highly influenced by its surroundings or whether the amino acid is on the N-terminal or not^37^.

To investigate how these interactions are distributed throughout the peptide, the comparison of amino acid contribution across retention mechanisms (RP and HILIC) for multiple peptides was plotted (**Fig. 3c**). The atomic saliency values were dataset-wise and peptide-wise normalized, and then they were summed for each amino acid. For this analysis, common peptides in both RP (SWATH Library) and HILIC (LUNA HILIC) datasets are considered. These peptides are aligned vertically by increasing molecular weight, and are displayed as mirrored sequences. In agreement with the single amino acid substitution experiments done by Kovacs et al.^38^, in RP, we observe the most significant positive contribution to retention for Phenylalanine (F) and Tryptophan (W). However, when these residues are positioned at the N-terminus, their hydrophobic contribution may be reduced—or even reversed—due to the presence of an uncompensated positive charge on the N-terminal amino group. We observe a repulsive effect at the C-terminus under RP conditions, whereas in HILIC, the C-terminal region contributes mainly positively to retention. In HILIC, the positive contribution of amino acids is distributed along the entire peptide sequence, with no significant deviation when located at the N-terminus.

The most notable difference between RP and HILIC retention behaviour in **Fig. 3c** is the overall shift towards negative contributions in RP, where peptides appear predominantly “red”. RP consists of a non-polar stationary phase, which interacts primarily through weak dispersion forces, and a polar mobile phase with strong polar interactions. Since polar interactions are stronger than hydrophobic ones, peptide retention in RP is primarily driven by the reduction of interactions between the analyte and the mobile phase. Consequently, peptides are retained when their affinity for the mobile phase decreases^39^. Gradient elution conditions with continuous change in the mobile phase elution power, therefore, represent the most critical parameters affecting retention in RP chromatography.

### Amino acid contributions on the dataset level

To examine the contributions of amino acids to retention, mean saliency values across ten datasets for individual amino acids are compared (**Fig. 4**). For this analysis, positional effects within the peptide were ignored and the saliency values were used without normalization. There is a decreasing positive contribution to retention from nonpolar to polar amino acids for RP chromatography. In contrast, HILIC exhibits the opposite trend, and the retention increases for polar amino acids. The SCX dataset shows the contribution to retention only for basic amino acids. Additionally, the decrease in repulsion with the increasing polarity of amino acid residues can be seen in SCX. To explore the combined effect of contribution and abundance of individual amino acids, summed saliency values were visualized, and the same trend was observed (data not shown). Moreover, **Fig. SI-9** confirms that even the dataset-wise normalization of mean saliency values for each amino acid provides the same trends.

**Figure 4.**
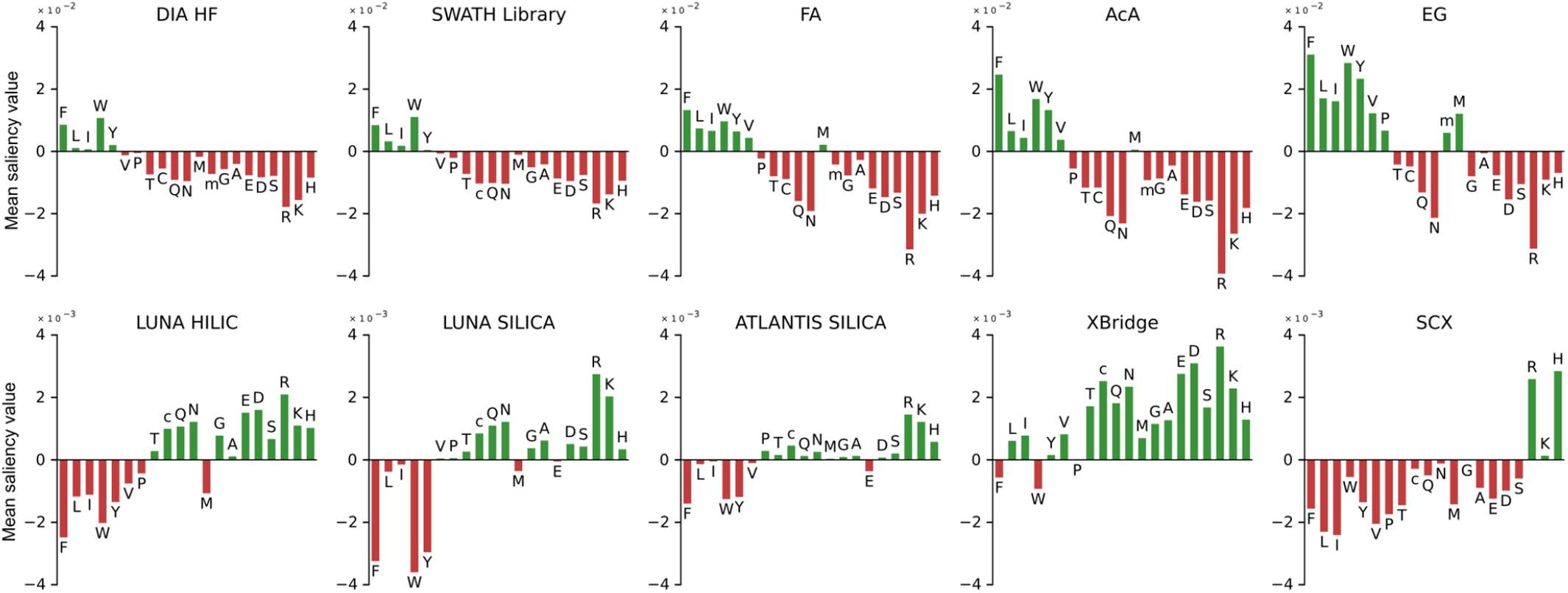
Amino acid contributions per dataset expressed as a mean saliency value for each amino acid. Saliency values were not normalized. M, methionine; m, oxidation of methionine; C, cysteine; c, carbamidomethylation of cysteine. Amino acids are ordered based on their decreasing hydrophobicity coefficients derived from RP experiments using linear regression analysis^40^.

### Effect of neighbouring amino acids

Peptide retention time is influenced by amino acid composition^2^, the position of each amino acid^41^, and peptide length^42^. Peptides with identical compositions but slight sequence variations exhibit different retention times, which can be explained by conformational effects and nearest-neighbor interactions^32^. To investigate these factors, we examined how specific residues affect retention along the peptide sequence.

**Fig. 5** shows heat maps of the mean saliency values for each amino acid (rows), computed from all instances where a given amino acid (columns) appears one position to the left (N-terminal side) for all investigated datasets. For completeness, the analogous analysis for the first position from the right (C-terminal side) is shown in Supplementary **Fig. SI-10**. Evaluating neighbouring amino acids for all datasets confirms the sequence-dependent nature of peptide retention across retention mechanisms (RP, HILIC, SCX). HILIC datasets show a larger sequence-retention dependency compared to RP and SCX. Even small RP datasets (with 2,000 peptides) provide more consistent saliency values per amino acid, regardless of their surroundings, compared to large HILIC datasets (with 40,000 peptides). The results also indicate that methionine plays a particular role, as it tends to slightly enhance the saliency values of surrounding amino acids in RP datasets. As methionine lacks strong hydrogen bonding or polar stabilization, surrounding residues are more exposed to the mobile phase, increasing their retention or repulsion. In terms of HILIC, on silica-based columns (ATLANTIS SILICA, LUNA SILICA), the retention is more influenced by its neighbouring residues than in bonded-phase HILIC columns (LUNA HILIC, XBridge). This is reflected by the larger variation in amino acid saliency values across individual rows of the heat maps, indicating that the contribution of an amino acid to retention depends more strongly on its peptide sequence. Supporting **Fig. SI-11** and **SI-12** illustrate how saliency values vary when amino acids occur at the first, fifth, and tenth positions on either the N- or C-terminal side of the studied residue, shown for both the RP (SWATH Library) and HILIC (ATLANTIS SILICA). The evaluation of both the N- and Cterminals reveals that the influence of a particular amino acid on neighboring residues decreases with distance.

**Figure 5.**
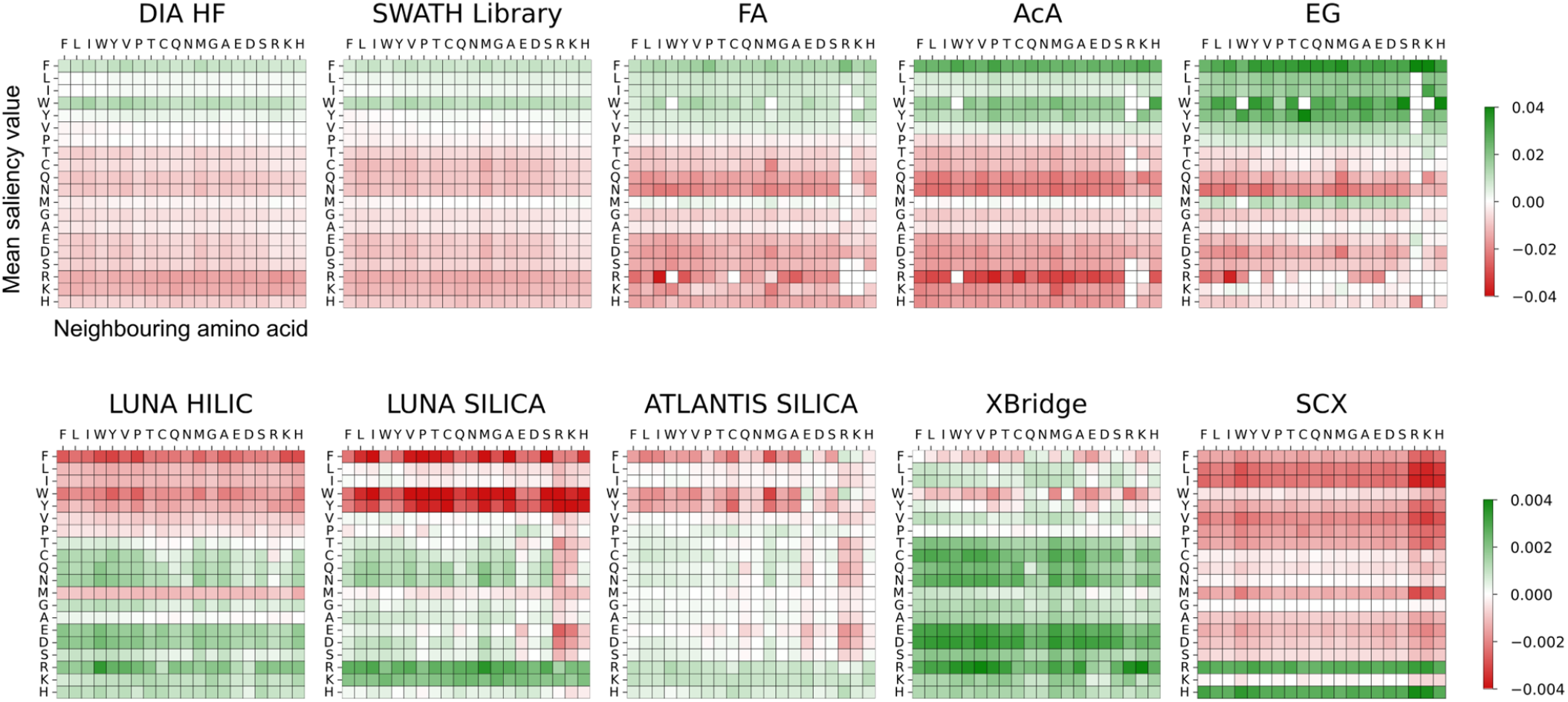
Heat maps for ten studied datasets of mean saliency values of an amino acid (rows) when a specific amino acid (columns) is present at the immediately adjacent N-terminal position. The colour scale indicates the magnitude of the retention (green) and repulsion (red). Saliency values were not normalized.

**Figure 6.**
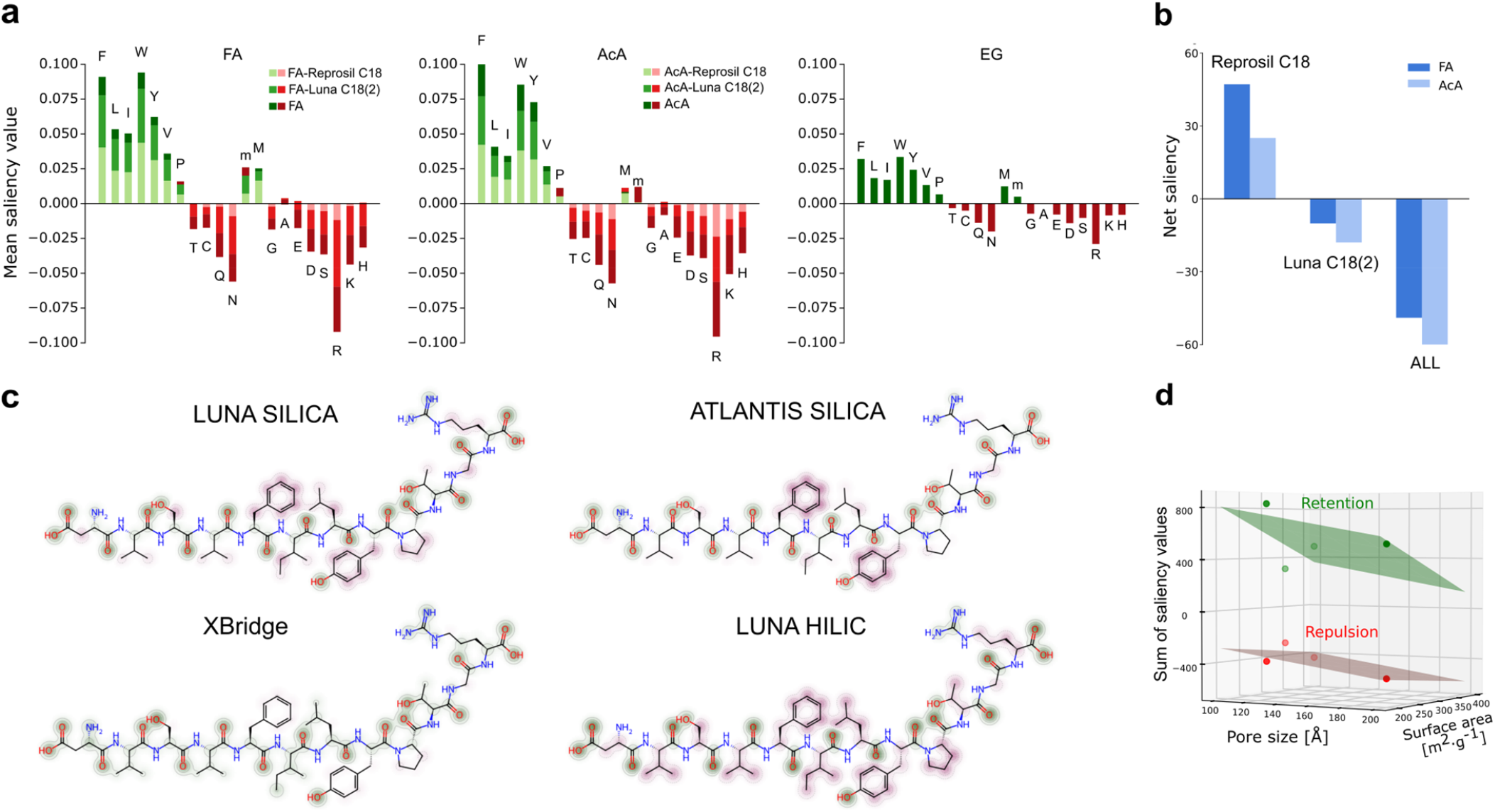
Saliency mapping for proteomics applications. (a) Mean amino acid saliency values compared for different chromatographic columns (Reprosil C18, Luna C18(2)) and mobile phase additives (FA, AcA, EG) in RP. M, methionine; m, oxidation of methionine; C, cysteine; c, carbamidomethylation of cysteine. (b) Bar plot of “net saliency” for Reprosil C18 and Luna C18(2) columns, and for both columns trained together (ALL). The net saliency refers to the sum of all saliency values. Saliency values were not normalized for this purpose. (c) Saliency maps for peptide DVSVFILYPTGR normalized on the dataset level and subsequently at the peptide level within the four HILIC datasets. (d) 3D scatter plot of the effect of pore size and surface area on retention (sum of all positive saliency values, green) and repulsion (sum of all negative saliency values, red) for HILIC columns.

### The effect of mobile phase additive in reversed-phase chromatography

Reversed-phase chromatography remains the most common method for final peptide separation prior to mass spectrometry. The mobile phase is typically an acidified acetonitrile-water binary mixture, which helps minimize the negative effects of residual silanol groups on silica columns, reduces metal-chelating tendencies of acidic peptides, and improves electrospray ionization efficiency^43^. In chromatography, the acidifier forms neutral ion pairs with protonated peptides, which modulate their retention in RP. The most common acidifier is 0.1% formic acid (FA), though some proteomics laboratories prefer 0.5% acetic acid (AcA), which achieves a comparable pH at that concentration^27^. Formic acid provides limited protonation in LC and does not enhance MS ionization. Therefore, mobile phases with FA are often supplemented with auxiliary additives such as dimethyl sulfoxide (DMSO) or heptafluorobutyric acid (HFBA). Recently, ethylene glycol (EG) has emerged as a protic alternative to DMSO, which, while effective, is known to cause carryover effects in MS instrumentation^44^.

These auxiliary additives enhance electrospray ionization, yet their effect on peptide retention has not been studied comprehensively. Therefore, datasets with different mobile phases and column chemistries were trained separately in PeptideGNN and visualized by saliency mapping. These datasets, which come from an interlaboratory study of two labs, differ solely in the mobile phase additive and chromatographic column. **Fig. 6a** shows stacked bar charts of the mean amino acid saliency values for RP datasets using three mobile phase additives (FA, AcA, EG) and two C18 columns (Reprosil C18, and Luna C18(2))^27^.

Bar charts were calculated for the same peptides identified across datasets. The overall retention trend for RP was consistent across all additives and chromatographic columns. However, subtle differences in the intrinsic contributions of individual amino acids were observed between FA and AcA. Because the mobile phase with EG was further acidified with 0.1% FA, the EG dataset mimics the behaviour of the dataset when only FA was used as the additive.

The sum of all saliency values within the dataset, including both positive and negative values, indicates the primary contribution to the retention mechanism in the chromatographic column. In chromatography, the analyte separation is governed by the balance between retention, driven by the affinity to the stationary phase, and repulsion, controlled by the affinity to the mobile phase. This difference between retention and repulsion is, in this case, referred to as “net saliency”. **Fig. 6b** shows net saliency values for FA and AcA datasets when models were trained separately for each column (Luna C18(2), Reprosil C18) or on datasets pooled across column chemistries, while preserving the identity of the mobile phase additive (ALL). When both column chemistries are trained together (ALL), AcA has a higher repulsive contribution compared to FA. The same trend was observed for the Luna C18(2) dataset, which constitutes 65% of the pooled ALL dataset. The trend differed for the Reprosil C18 dataset, where overall retention seemed dependent on retention rather than repulsion. Net saliencies confirm previously published results that ion-pairing with AcA provides more hydrophilic ion pairs compared to FA, resulting in lower peptide retention when using AcA^27^. For ALL and Luna C18(2) datasets, stronger repulsion for AcA was observed (negative values of “net saliency”), which corresponds to the phenomenon described earlier, where the main force controlling retention in RP is the enhanced repulsion (**Fig. 3c**). In the Reprosil C18 dataset, the less hydrophobic Reprosil-PUR C18-AQ column was used, which provided separation even of hydrophilic peptides, suggesting that the main force is retention over repulsion (as demonstrated by the positive values of “net saliency” in **Fig. 6b**). In addition, Reprosil-PUR C18-AQ has a higher surface area than Luna C18(2), which also leads to higher retention. The results shown in **Fig. 6a** and **Fig. 6b** demonstrate the ability of PeptideGNN to recognize the subtle interaction differences caused by the type of mobile phase additive or chromatographic column, and to report on these differences through saliency mapping.

### Characterization of HILIC columns using saliency mapping

Despite its popularity, RP has notable drawbacks, including peptide overloading^45^ and peak distortions^46^. To overcome these problems, chromatographic columns with polar stationary phases performing HILIC mechanisms have been adopted in proteomics^29^. HILIC utilizes well-established silica-based columns, which can be modified with various bonded phases, making the selection of an appropriate stationary phase for a specific proteomic application particularly challenging. Here, we establish the use of saliency maps to compare four different HILIC stationary phases. These include two bonded-phase columns - one functionalized with diol groups (LUNA HILIC) and the other with amide groups (XBridge) - as well as two unmodified bare silica stationary phases (ATLANTIS SILICA and LUNA SILICA). Retention times for the HILIC datasets were used as provided, including the data processing performed by the original authors according to the previously published protocol^47^. Supplementary **Fig. SI-13** shows model performance and saliency mapping results when investigating raw HILIC contribution, where the fraction numbers were used as the output for the PeptideGNN model. We observed the same trends in saliency mapping when predicting fractions instead of retention times, although some subtle nuances were lost due to the coarser granularity of fraction-based data.

In **Fig. 4**, the amino acid contributions for all four HILIC datasets are shown. For clarity, Supplementary **Fig. SI-14** presents bar charts only for the identical peptides identified across all four HILIC datasets. Both visualizations reveal the same overall trends, observable at both the amino acid level and the atomic level when examining saliency maps of individual peptides. For demonstration, the peptide DVSVFILYPTGR was selected because it was identified in all four datasets (**Fig. 6c**). The saliency maps for this peptide were normalized both dataset-wise and peptide-wise to ensure full comparability in terms of interaction patterns and saliency intensities. On bare silica stationary phases (LUNA SILICA and ATLANTIS SILICA), positively charged residues (arginine, lysine, histidine) exhibit strong retention by hydrogen bonding and ion-exchange interactions with silanols (Si-OH) on stationary phase, while other residues do not contribute significantly to the retention (**Fig. 4**). In contrast, the retention and repulsion is spread over all amino acids for bonded stationary phases (XBridge, LUNA HILIC), suggesting that they are more targeted for their use in proteomics. The amide-bonded stationary phase (XBridge) shows higher retention for most amino acid residues (**Fig. 4, Fig. 6c**), consistent with its established adsorptive retention mechanism^48^. Unlike bare silica, amide columns do not exhibit ion-exchange mechanism^39^; instead, retention is governed primarily by hydrogen bonding and dipole-dipole interactions. These interactions target a broader range of amino-acid properties. Consequently, the XBridge amide phase is particularly suitable for separating highly hydrophilic analytes, such as oligosaccharides or glycopeptides, which cannot be distinguished using ion-exchange mechanism of bare silica. Stationary phases with chemically bonded diol groups (LUNA HILIC) retain analytes primarily through partitioning between the aqueous layer on the stationary phase and the organic-rich mobile phase^49^. Due to that, its retention pattern is inversely proportional to the RP mechanism (**Fig. 4**), where the leading mechanism is also partitioning. Notably, normalized saliency maps (**Fig. 6c**) show higher saliency values for LUNA SILICA compared to ATLANTIS SILICA. Since both datasets were measured under identical type of stationary phase surface and experimental conditions within one lab, the observed differences can be attributed to the hydrodynamic properties of the stationary phases. We therefore suggest that the stronger retention on LUNA SILICA arises from its higher surface area, which provides more interaction sides on the silica surface.

To study this effect, we visualized how peptide interaction (green) and repulsion (red) vary with the physico-chemical properties of HILIC stationary phases, such as pore size and surface area by looking at the sum of all positive and negative saliency values (**Fig. 6d**). The highest peptide retention is provided by the combination of the small pore size and surface area, while the column with the larger values of pore size and surface area shows the highest repulsion. Higher retention effect of columns with smaller pore sizes is probably related to the minor difference between the pore size and the hydrodynamic diameter of the peptide^44^. In case of columns with a significant repulsion (high surface area, large pore size), the high surface area contribution suggests active role of the stationary phase to the repulsion, i.e., the opposite effect to RP, where the repulsion is mainly controlled by the mobile phase composition.

Nevertheless, the dominant effect of the stationary-phase chemistry and mobile phase composition on peptide retention cannot be omitted. Hence, **Fig. 6d** illustrates an additional feature of the PeptideGNN model, allowing a better understanding of peptide interactions in the HILIC chromatography.

## Conclusion

Apart to strong predictive performance and generalizability to non-canonical amino acids, the presented PeptideGNN model can also be explained by using a saliency mapping method, demonstrating it had learned the underlying principles of all major peptide retention mechanisms directly from sequence and retention time alone. This illustrates its ability to serve as an optimisation tool in the analysis and design of various chromatographic experiments.

Indeed, the straightforward visualization and analysis of PeptideGNN-derived saliency maps provides a unique tool to improve our understanding of the most common retention mechanisms, and to extend the detailed description of peptide-stationary phase interactions. This is illustrated by PeptideGNN’s unprecedented ability to distinguish subtle interaction differences arising from various chromatographic parameters, including stationary phase chemistry and the type of mobile phase additive.

Saliency maps at amino acid resolution revealed substantial contributions of neighboring residues to peptide retention. These findings provide a background for the rational design of proteomics workflows, enabling improved peptide selection for targeted assays and more effective control over chromatographic separation, particularly for challenging sequence motifs and post-translationally modified peptides.

As the HILIC retention mechanism offers an interesting alternative in peptide separation, we investigated the impact of the stationary phase surface chemistry on HILIC column retention. The results of saliency mapping revealed that retention on bare silica columns is primarily driven by interactions with positively charged amino acid residues. In contrast, bonded stationary phases interact with all amino acids of the peptide, which makes them more suitable for proteomics applications.

Overall, the resulting patterns of saliency maps reflect established chromatographic theory and are consistent with previously published results, providing strong evidence that PeptideGNN provides a unique means to deliver a highly performant (modified) peptide retention predictor, but also to reveal the physicochemical logic governing peptide separation.

## Data availability

The raw mass spectrometry data used in this study are publicly available via the PRIDE repository. A complete list of raw data identifiers and corresponding publication DOIs are provided in Supporting information and in the README file of the PeptideGNN GitHub repository. See the repository at https://github.com/CompOmics/peptide-gnn.

The processed datasets generated after open-search and used for training and evaluation of the PeptideGNN model are also available in the GitHub repository.

## Code availability

For the implementation, training and interpretation of PeptideGNN see https://github.com/CompOmics/peptide-gnn.

## Acknowledgements

A.K., R.D, L.M. and R.B. acknowledge funding from the Research Foundation Flanders (FWO) [G010023N, 1SH9O24N, G0GDV23N, 12A6L24N]. L.M. acknowledges funding from the Horizon Europe Projects BAXERNA 2.0 [101080544] and COMBINE [101191739], and from the Ghent University Concerted Research Action [BOF21/GOA/033]. K.H. acknowledges Brno Ph.D. Talent Scholarship – Funded by the Brno City Municipality. The financial support from the Czech Science Foundation, project 23–07581S, is gratefully acknowledged by K.H. and J.U.

## Author contributions

A.K. conceptualized and developed the PeptideGNN model, K.H. conceptualized and performed the experiments and analyses, contributed to the model development, R.D., A.N., A.D., and R.G. revised the manuscript, A.K., K.H., R.B., and J.U. wrote the manuscript, L.M., R.B., and J.U. conceptualized and supervised the study, and revised the manuscript. A.K. and K.H. contributed equally to this work.

## Correspondence and requests for materials

The requests should be addressed to Robbin Bouwmeester or Jiri Urban.

## Competing interests

The authors declare no competing interests.

## Additional information

Supplementary information is attached as individual file.

## Notes

https://github.com/CompOmics/peptide-gnn

### Competing Interest Statement

The authors have declared no competing interest.

