## Supplementary Table 1; Supplementary Figures 1 - 14 for "Predicting and Elucidating Peptide Retention Mechanisms with Graph Attention Networks"

***Table SI-1.*** *Overview of data sets used with detailed information about their chromatographic conditions.*

| Dataset | Column | ID x L, mm | Chemistry | $D_p$, µm | Surface area, m$^2$/g | Pore size, Å | Mobile phase | Fractionation | Source |
|---|---|---|---|---|---|---|---|---|---|
| DIA HF | ReproSil-Pur C18-AQ | 0.075 x 500 | C18, endcapping | 1.9 | 300 | 120 | Water / ACN / FA | High pHRP (0. Reversed- M AmF) and SAX | MCP 16: 10.1074/mcp.RA117.000314, 2296–2309, 2017. |
| SWATH Library | Magic (Bischoff Chromatograph y) | 0.075 x 200 | C18 | 3 | - | 200 | Water / ACN / FA | SAX | Sci Data 1: 10.1038/sdata.2014.31 |
| FA, AcA (UM) | Luna<br>C18 | 0.100 x 200<br>0.100 x 150 | C18 | 3 | 400 | 100 | Water / ACN / FA / AcA | -, High pH RP | J. Proteome Res. 2023, 22, 272–278 |
| FA, AcA (KU) | ReproSil-Pur C18-AQ | 0.075 x 500 | C18, endcapping | 1.9 | 300 | 120 | Water / ACN / FA / AcA | -, High pH RP | J. Proteome Res. 2023, 22, 272–278 |
| EG | Reprosil Gold C18 | 0.075 x 400 | C18 | 3 | 300 | 100 | Water / ACN / FA / EG | - | Anal Bioanal Chem (2017) 409:1049–1057 |
| Luna Silica | Luna Silica | 3.0 × 50 | Unbonded silica | 3 | 400 | 100 | 10 mM ammo-nium formate pH 4.5 / ACN | (HILIC x RP, Luna C18(2)) | JCA, 1534 (2018) 75–84 |
| Luna HILIC | Luna HILIC | 3.0 × 50 | Diol, Ethylene bridges | 3 | 200 | 200 | 10 mM ammo-nium formate pH 4.5 / ACN | (HILIC x RP, Luna C18(2)) | JCA, 1534 (2018) 75–84 |
| Atlantis SIlica | Atlantis Silica | 3.0 × 50 | Silica | 3 | 330 | 100 | 10 mM ammo-nium formate pH 4.5 / ACN | (HILIC x RP, Luna C18(2)) | JCA, 1534 (2018) 75–84 |
| XBridge | Xbridge Amide | 3.0 × 50 | Amide (-linker-CONH2) | 3.5 | 185 | 130 | 10 mM ammo-nium formate pH 4.5 / ACN | (HILIC x RP, Luna C18(2)) | JCA, 1534 (2018) 75–84 |
| SCX | Polysulfoethyl A | 2.1 x 100 | poly(2-sulfoethyl aspartamide) | 5 | - | 200 | Water / ACN / 500 mM KCl | (SCX x RP, Luna C18(2)) | Anal. Chem. 2017, 89, 11795-11802 |

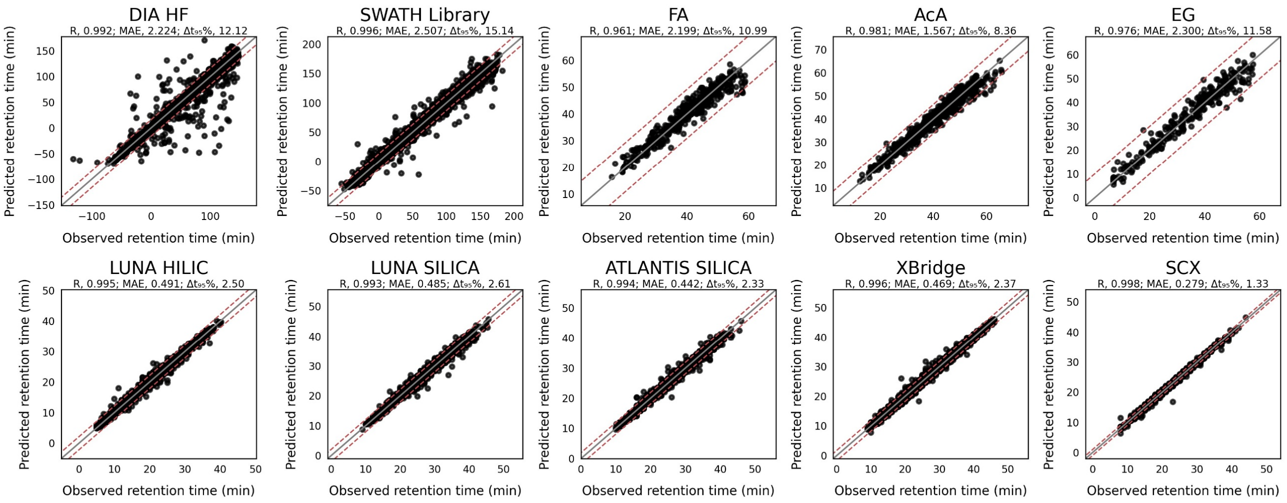


***Figure SI-1.*** *Scatter plots of predicted against observed retention times on the test data of ten studied datasets. The red dotted line indicates the 99th error percentile.*

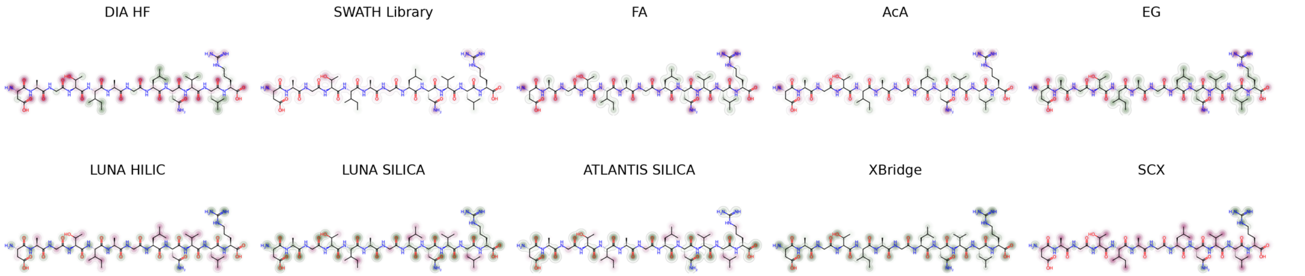


***Figure SI-2.*** *Saliency maps for the peptide DAGTIAGLNVLR generated from all ten trained datasets.*

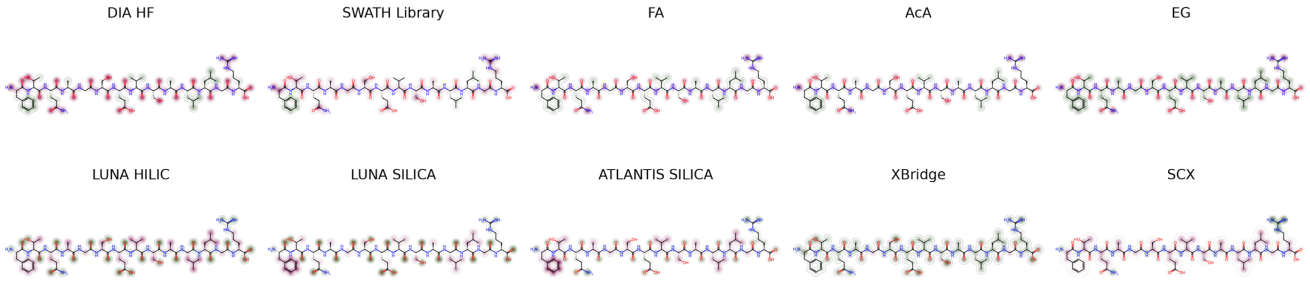


***Figure SI-3.*** *Saliency maps for the peptide FTQAGSEVSALLGR generated from all ten trained datasets.*

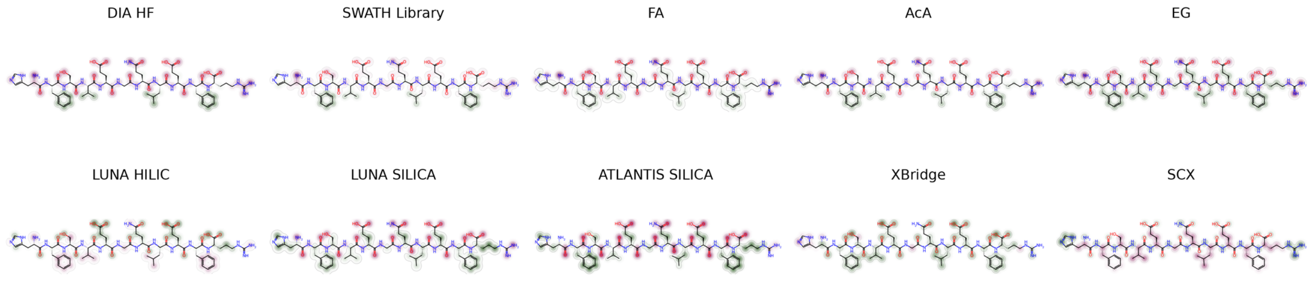


***Figure SI-4.*** *Saliency maps for the peptide HFSVEGQLEFR generated from all ten trained datasets.*

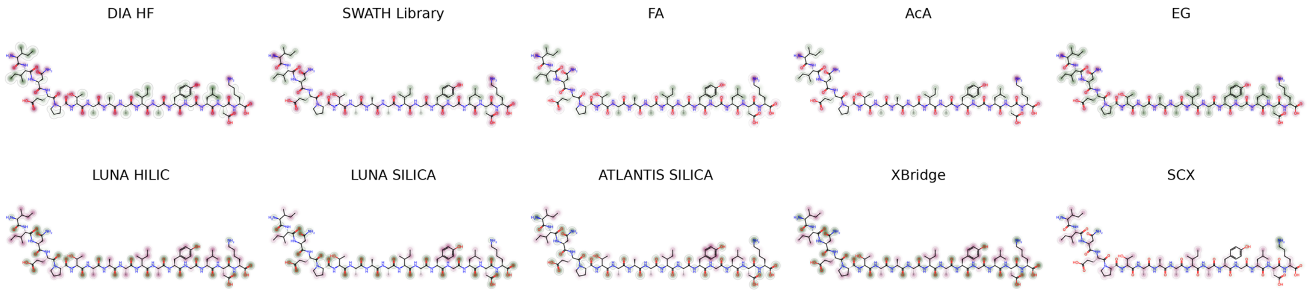


***Figure SI-5.*** *Saliency maps for the peptide IINEPTAAAIAYGLDK generated from all ten trained datasets.*

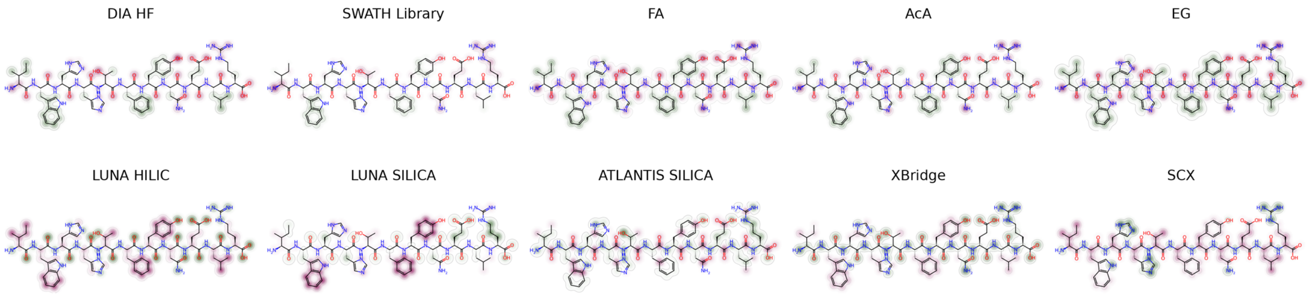


***Figure SI-6.*** *Saliency maps for the peptide IWHHTFYNELR generated from all ten trained datasets.*

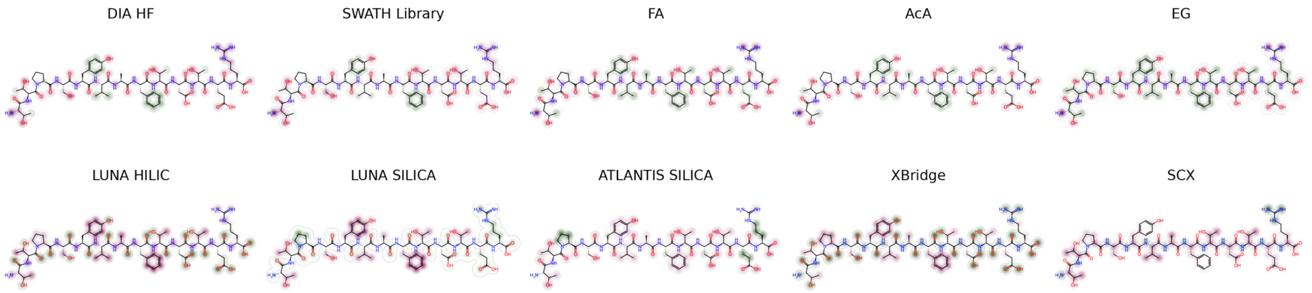


***Figure SI-7.*** *Saliency maps for the peptide TTPSYVAFTDTER generated from all ten trained datasets.*

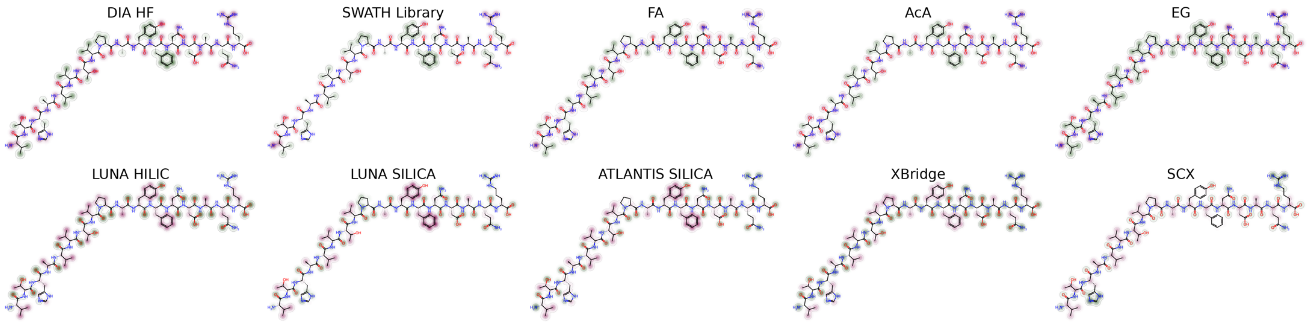


***Figure SI-8.*** *Saliency maps for the peptide VTHAVVTVPAYFNDAQR generated from all ten trained datasets.*

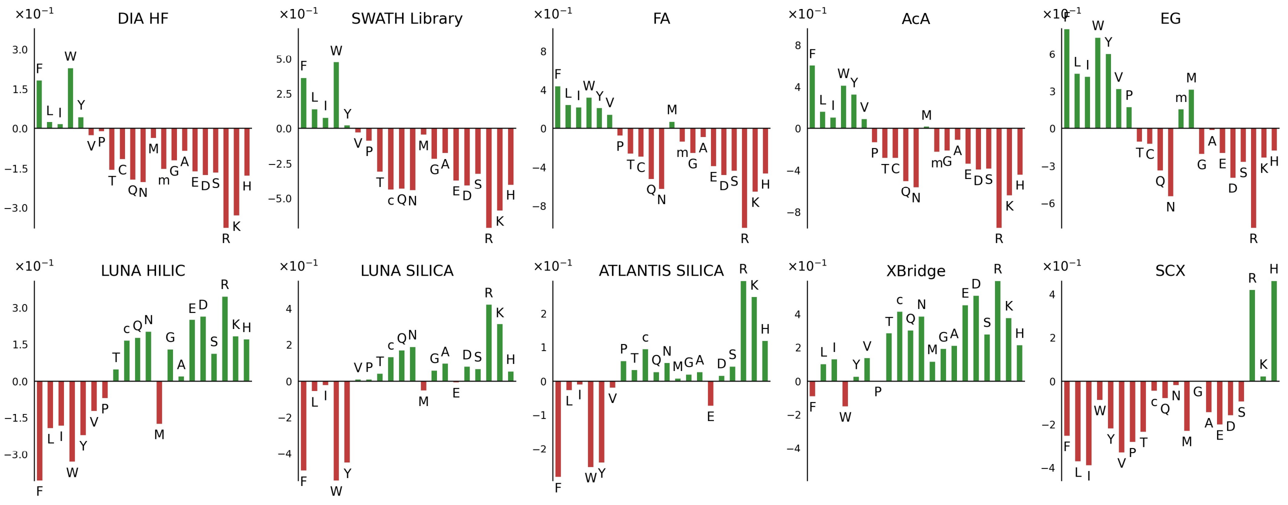


***Figure SI-9.*** *The bar charts of amino acid contributions per dataset are expressed as a mean saliency value for each amino acid. Saliency values were dataset-wise normalized by dividing atomic saliency values by the maximal absolute saliency value found within the dataset. M, methionine; m, oxidation of methionine; C, cysteine; c, carbamidomethylation of cysteine.*

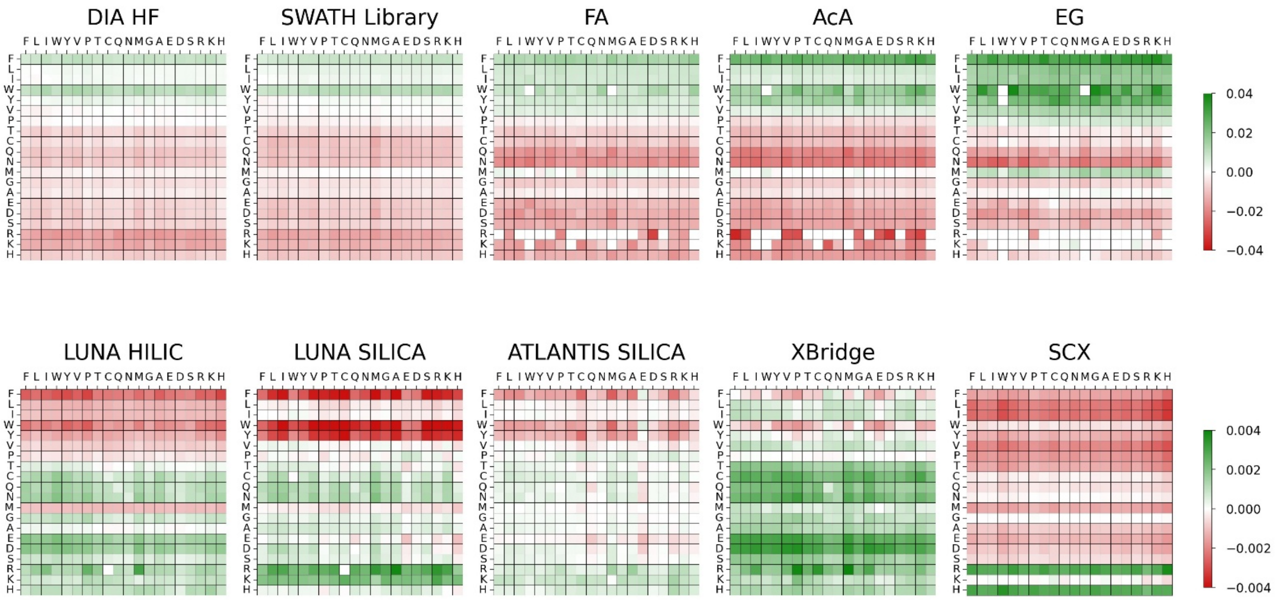


***Figure SI-10.*** *Heat maps for ten studied datasets of mean saliency values of an amino acid (rows) when a specific amino acid (columns) is present at the first position from the C-terminal. The colour scale indicates the magnitude of the interaction (green) and repulsion (red). Saliency values were not normalized.*

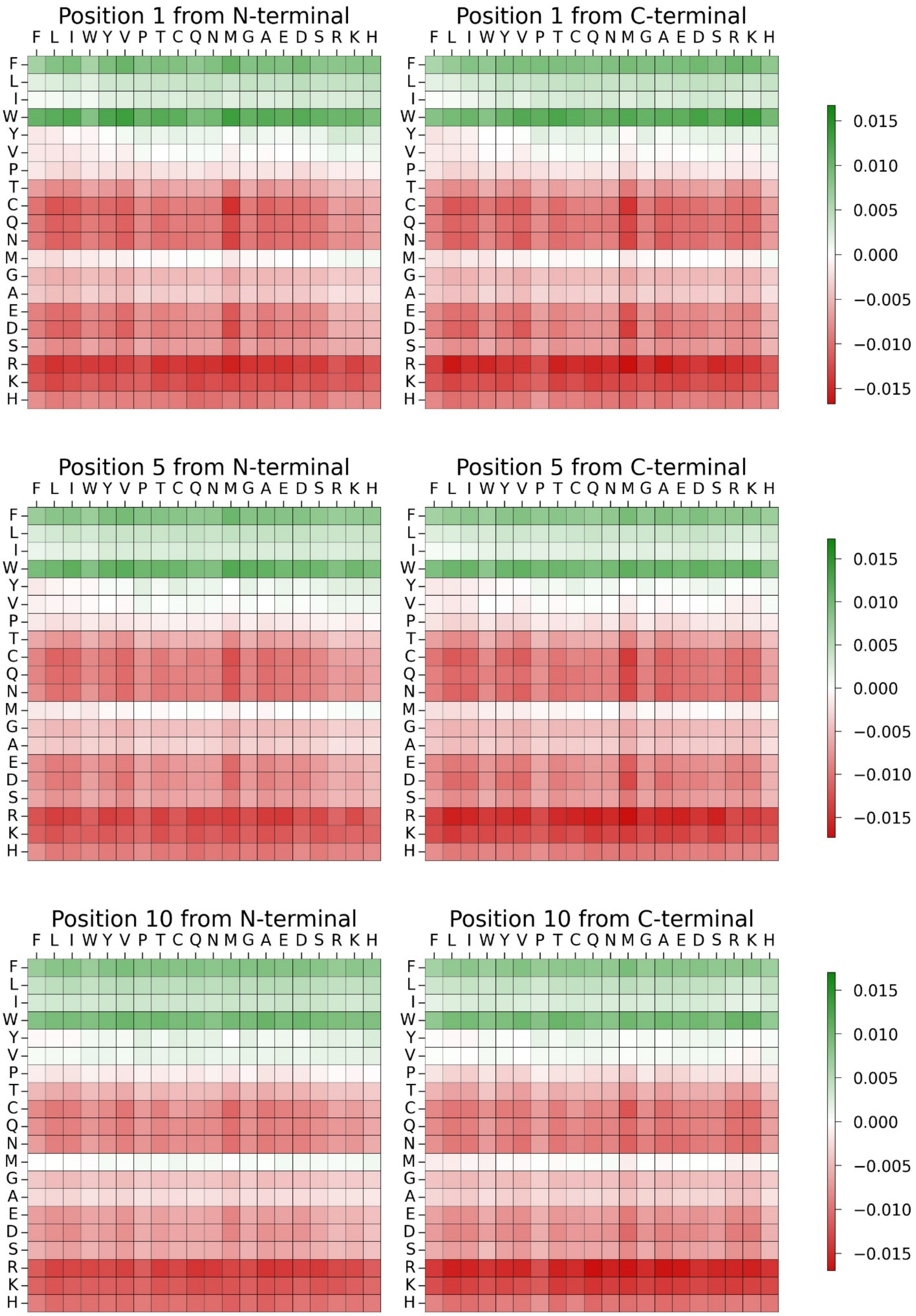


***Figure SI-11.*** *Six heat maps for SWATH Library data set of mean saliency values of an amino acid (rows) when a specific amino acid (columns) is present at the 1st, 5th, and 10th position from N-terminal and C-terminal. Saliency values were not normalized.*

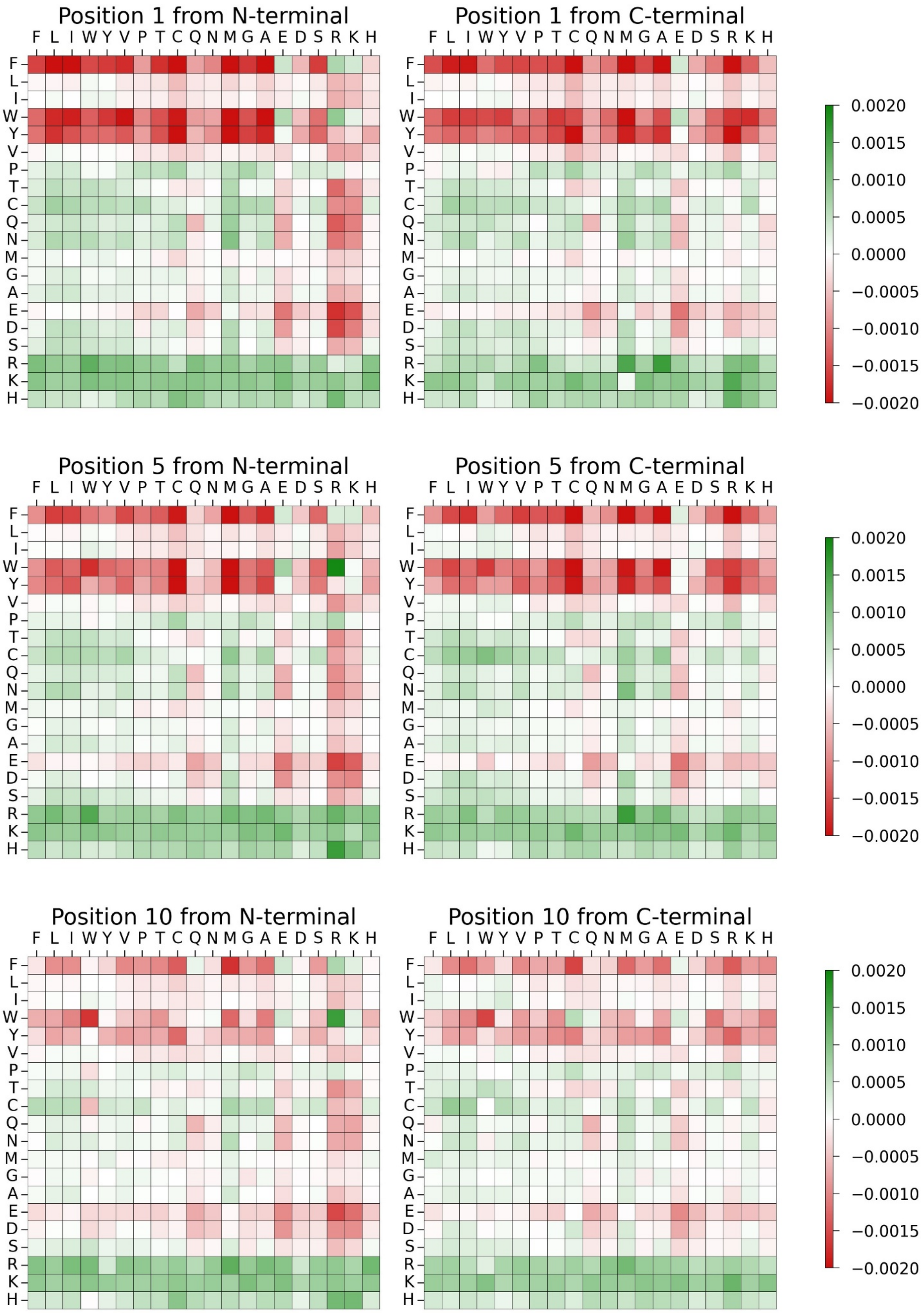


***Figure SI-12.*** *Six heat maps for ATLANTIS SILICA data set of mean saliency values of an amino acid (rows) when a specific amino acid (columns) is present at the 1st, 5th, and 10th position from N-terminal and C-terminal. Saliency values were not normalized.*

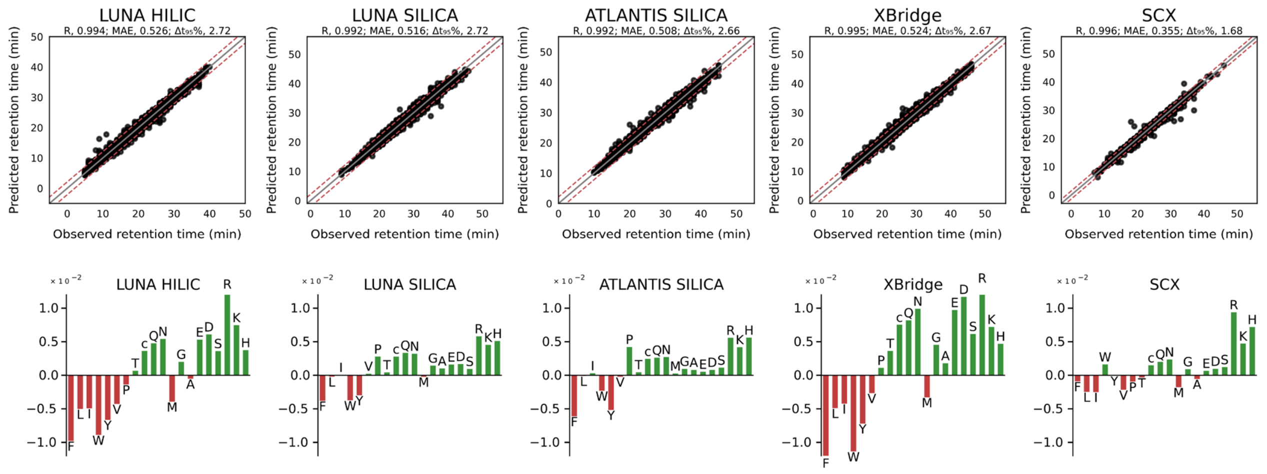


***Figure SI-13.*** *Scatter plots for PeptideGNN model performance for predicting fraction numbers for HILIC and SCX datasets. Bar charts of mean saliency values were computed from the trained model. Amino acids in the bar charts were ordered by their increasing polarity. Saliency values were not normalized.*

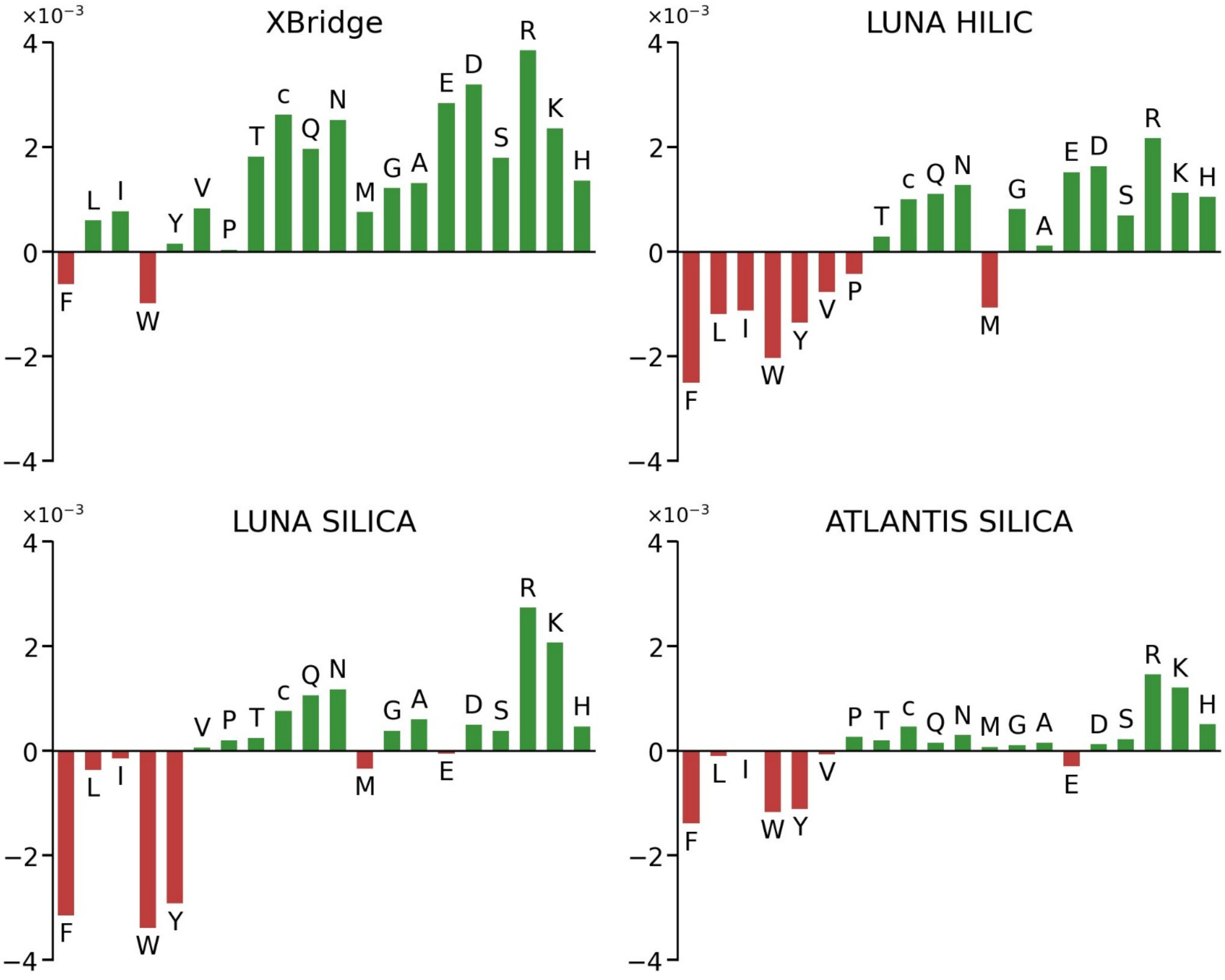


***Figure SI-14.*** *The bar charts of amino acid contributions for HILIC data sets expressed as a mean saliency value for each amino acid. Post-translational modifications are signed with lowercase letters. Saliency values were not normalized; bar charts were plotted on 21063 common peptides found in all datasets.*
